# Drone methods and educational resources for plant science and agriculture

**DOI:** 10.1101/2025.05.19.654604

**Authors:** Travis Parker, Burcu Celebioglu, Mark Watson, Paul Gepts

## Abstract

Technological advances have made drones (UAVs) increasingly important tools for the collection of trait data in plant science. Many costs for the analysis of plant populations have dropped precipitously in recent decades, particularly for genetic sequencing. Similarly, hardware advances have made it increasingly simple and practical to capture drone imagery of plant populations.However, converting this imagery into high-precision and high-throughput tabular data has become a major bottleneck in plant science. Here, we describe high-throughput phenotyping methods for the analysis of numerous plant traits based on imagery from diverse sensor types. Methods can be flexibly combined to extract data related to canopy temperature, area, height, volume, vegetation indices, and summary statistics derived from complex segmentations and classifications. We then describe educational and training resources for these methods, including a web page (PlantScienceDroneMethods.github.io) and an educational YouTube channel (https://www.youtube.com/@travisparkerplantscience) with step-by-step protocols, example data, and example scripts for the whole drone data processing pipeline. These resources facilitate the extraction of high-throughput and high-precision phenomic data, removing barriers to the phenomic analysis of large plant populations.

## 1. Introduction

Global demand for agricultural products is projected to increase by 50-60 percent between 2019 and 2050 (Falcon et al., 2022), with even greater required increases in many regions. For example, in Sub-Saharan Africa, demand for cereal grains and other crops is expected to triple between 2010 and 2050 (Van Ittersum et al., 2016). Current trajectories are insufficient to achieve sustainable development goals (FAO, 2018; Van Ittersum et al., 2016). Achieving the required increases in crop productivity will require significant gains in breeding and improved crop management. These increases will require evaluations of large plant populations and the generation of high-precision, high-throughput data on an unprecedented scale.

The pace of technological advances in other areas, such as DNA sequencing and genomics, has greatly outpaced advances in phenotyping in recent years. This has left trait data collection as a major remaining bottleneck in many plant science studies. Simultaneously, where technological advances in phenotyping have occurred, the data processing pipelines required to effectively apply these new technologies are often lacking. Development of efficient pipelines and training materials for high-throughput and high-precision phenotyping are of foremost value in biological sciences.

Drones, also known as Unoccupied Aerial Vehicles (UAVs), or sometimes Uncrewed or Unmanned Aerial Vehicles, are the central component of Unoccupied Aerial Systems (UAS), and have undergone rapid technological advances in the last decade (Rejeb et al., 2022; Sankaran et al., 2015; Pungavi and Praveenkumar, 2024; Xie and Yang, 2020; Yang et al., 2017; Pathak et al., 2022). Increasing affordability and practicality of drone use in plant sciences has been made possible by improvements and miniaturization of essential components such as microprocessors, cameras, inertial measuring units, and batteries. The quality and price of these technologies has improved due to their broad use in mass-produced consumer products such as smartphones and tablets.

Unlike traditional phenotyping strategies, the cost and time investment in drone-based phenotyping does not scale linearly with the number of sampled units, such as individual plants or plots. Instead, data can be mass-extracted from large numbers of experimental units with little additional time or cost invested by increasing sample size. This is particularly true when processing pipelines can be automated. This scaling makes drones a particularly valuable tool for high-throughput phenotyping (HTP) of large populations, such as those required in Genome Wide Association Studies (GWAS), Quantitative Trait Locus (QTL) mapping, plant breeding, phenomic selection, or surveying of natural populations (Xiao et al. 2022). Additionally, drones offer advantages in precision and flexibility relative to other approaches. The spatial resolution (ground sampling distance, GSD) of drone imagery is typically approximately 0.5-20 cm px^-1^, considerably higher than the >100 cm resolution typically achieved by satellites or crewed aircraft (Inoue, 2020; Bansod et al., 2017). This high level of detail is required for many studies of plant canopy health and growth. Drones also offer flexibility in temporal resolution relative to other aerial imagery options and can be operated below cloud cover. Compared to ground-based imaging platforms, drones are also typically compact, inexpensive, and quick to deploy.

Software advances have also occurred in recent decades. In particular, open source software such as QGIS (QGIS development team, 2024) offer numerous functionalities for extraction of geographic data. QGIS is free and open-source, and runs on Windows, iOS, or Linux systems. This makes it readily accessible to research teams globally, including those on limited budgets and in the Global South. Additionally, advances in mission planning applications for automated drone flight, as well as structure-from-motion photogrammetry tools, have also simplified drone data collection and processing.

Despite these technological advances, the linking of each component in the processing pipeline and training teams on their use have often become limiting steps in application of drones in plant science. Here, we describe workflows for the generation and extraction of high-throughput and high-precision phenotype data from drone imagery. We then present free educational resources to train global teams on how to apply these methods to address diverse challenges. These resources will be important for resolving critical bottlenecks to the application of this promising emerging technology.

## 2. Materials and Methods

### 2.1 Imagery acquisition and mission planning applications

Full protocols: https://plantsciencedronemethods.github.io/pages/1.Introduction/Methods.html https://plantsciencedronemethods.github.io/pages/MissionPlanning.pdf

The methods described herein are compatible with a variety of aircraft and sensor types, and are optimized for use with multispectral, thermal, and RGB cameras. Mission-planning applications are important for collecting drone imagery at a uniform, desired spacing. Numerous simple and user-friendly applications exist for mission planning, which vary in compatibility with aircraft, cameras, and control devices. Major mission planning applications include DJI GS Pro, Pix4Dmapper, Drone Deploy, Map Pilot Pro, and UgCS, among numerous others. The basic premise of these is generally similar, with the user specifying flight boundaries, altitude, image overlap, and camera angle. For flights requiring multiple batteries, pause-and-resume features are available on most major applications, allowing for seamless battery swaps during missions.

For most camera types and applications, 70-85% side-and front-overlap between adjacent images is ideal. Higher overlap is important when mapping difficult areas, such as those with a high percentage of water or high visual uniformity. Wind and cloud cover, particularly partly cloudy conditions, are problematic for photogrammetry and should be avoided where possible.

In many cases, multispectral cameras are equipped with their own Global Positioning Satellite (GPS) unit and are triggered independently of the drone’s recognized camera, with the front-overlap (frontlap) triggering programmed separately. These cameras also often include downwelling light sensors to account for ambient lighting conditions. Calibration images of a target with known reflectance are optimal to correct for lighting conditions. Example cameras like this include the Micasense RedEdge and Altum series. In some cases, manually setting white balance and ISO settings on RGB cameras can improve image consistency and limit artifacts based on different lighting conditions across the field.

When data is captured over multiple flight dates, or with multiple cameras, alignment of raster outputs can greatly simplify the extraction of data from the files. This precise alignment can easily be achieved using ground control points (GCPs). Placing GCPs throughout the field is therefore strongly recommended. For most applications in the plant sciences, these do not need to be georeferenced with an additional Global Navigation Satellite System (GNSS; e.g., GPS) receiver.

The GNSS receivers on most commercial drones are accurate to 1-5 meters, and orthomosaics, models, and other photogrammetric outputs can be aligned to each other in relative space with centimeter-level resolution based on the GCPs in common between image sets. If very high precision alignment in absolute space is required, real-time kinematic (RTK) and post-processing kinematic (PPK) drones can be used to achieve calibrated GNSS spatial data with 1 cm scale precision.

Any object in a fixed (stationary) position and identifiable image sets or rasters can be used as a GCP. GCP tiles are commercially available or can be custom-made at low cost. In some cases, objects already located in the field (e.g., irrigation headers) can also be used as ground control points. Greater numbers of ground control points improve georeferencing quality, with fifteen or more ground control points generally being optimal (Agüera-Vega et al., 2017), while fewer may produce sufficient quality for many projects (Tahar, 2013; Oniga et al., 2018). Best practices for GCP placement include covering the full surveyed area, especially near the periphery, avoiding straight lines, and reasonably covering the full region of interest (Martínez-Carricondo et al., 2018).

Most jurisdictions have laws covering the use of drones, and these vary considerably by region. A comprehensive summary of these can be found at droneregulations.info (Accessed May 15, 2025).

Common restrictions include certification requirements to serve as remote pilot-in-command, limitations on airspace use, and specifications on the aircraft and hardware that can be flown, including distinctions based on manufacturer. Many institutions also have policy requirements on the use of drones in a workplace environment. It is the responsibility of the remote pilot-in-command to ensure that drone missions are conducted in compliance with local legislation and institutional requirements.

### 2.2. Structure from motion photogrammetry

Full Protocol (Pix4D): https://plantsciencedronemethods.github.io/pages/3.Pix4D/Methods.html.

After collecting raw imagery, the files are transferred off the camera’s memory card. Photogrammetry can be conducted using a variety of software options, including options that are proprietary (e.g., Pix4Dmapper, Agisoft Metashape, Drone Deploy, and Maps Made Easy), or free and open source (OpenDroneMap). While there is a cost to use commercial software, these may also provide simplicity, user-friendly and advanced features, fast turnaround times, and very high-quality outputs. Proprietary photogrammetry software options offer variable pricing structures and typically have free trial options. Open Drone Map is free to use, making it an attractive option for some research programs, although the installation is more complex and it may be comparatively limited in features.

Photogrammetric processing can be conducted on the cloud or on a local machine. Cloud-based processing often hastens processing times and frees up local computing resources, but in some cases, particularly in developing countries, image upload speeds can be limiting. Some institutions may also have requirements related to upload of sensitive data to cloud services. In these cases, processing on a local machine may be a more favorable option.

At a minimum, photogrammetric processing involves selection of raw images for processing, and the selection of the desired outputs. Photogrammetry software identifies common tie points between overlapping features in the image sets, and is able to calculate the geometry between these in a process called structure-from-motion, yielding 3-dimensional outputs from 2-D inputs (Ullman, 1979; Schonberger et al., 2016; Iglhaut et al., 2019). These outputs include orthomosaics, vegetation index maps, digital surface models (DSMs), point clouds of ties, and quality metrics. Optional photogrammetry steps include the addition of calibration images, particularly for multispectral cameras, and the addition of GCPs, which can also be added later in geographic information systems. The outputs of photogrammetry contain information related to plant canopy geometry (size, shape), plant number, and spectral reflectance data that can then be extracted in geographic information systems.

### 2.3. QGIS setup and georeferencing

Full protocols: https://plantsciencedronemethods.github.io/pages/4.QGIS_introduction/Methods.html https://plantsciencedronemethods.github.io/pages/7.Georeferencing/Methods.html

Drone-derived spatial data can be analyzed using Geographic Information Systems such as QGIS (formerly Quantum GIS, QGIS.org 2025) or ESRI ArcGIS. Here, we will focus on QGIS due to its free and open source model, which makes it broadly accessible globally. Similar methods are available in ArcGIS and other platforms.

QGIS is available for download at https://qgis.org/download/. Outputs from photogrammetry such as DSMs, vegetation indices, and other orthomosaics can be directly imported into a new project, and are represented in the Layers Panel, with the top files in the Layer Panel superimposed on the others in the map canvas.

Raster outputs from photogrammetry (e.g., reflectance maps, orthomosaics, point clouds, and DSMs) typically come with geospatial position data from the camera’s geotags. However, these are typically accurate only to within several (∼5) meters. Artifacts and distortion can also be introduced during photogrammetry. Photogrammetric outputs from different photogrammetric projects (e.g., from different cameras or different flight dates) are typically imprecisely aligned, often deviating by several meters without georeferencing.

Fortunately, these outputs can be easily aligned using QGIS’ Georeferencer functionality. To do this, the image sets must have visible features in common, called GCPs. In the QGIS georeferencer, the file to be georeferenced is loaded and GCPs are selected, and then the corresponding positions in the other raster file (being aligned to) can be selected from the map canvas. In the georeferencer, the “Thin Plate Spline” transformation algorithm may be particularly useful, as it is able to correct for warping and complex distortions in the map. This requires at least 10 GCPs. In contrast, the default transformation option (“Linear”) only changes the position and scale of the raster, without correcting for distortion. The “Cubic (4×4 kernel)” resampling method is well-suited for use with the Thin Plate Spline transformation algorithm. The georeferencer can then be run and outputs can be compared by selecting and deselecting in the Layers Panel to confirm alignment.

### 2.4. Setting up plot grids

Full protocol: https://plantsciencedronemethods.github.io/pages/5.PlotGrids/Methods.html

The “Create Grid” function in QGIS is well-suited to quickly establish a vector layer that distinguishes individual sampling units, such as plots, plants, trees, or quadrats. If plots are oriented at an angle geographically (i.e., not north to south, east to west), then the “Rotation” option for the map canvas can be used to simplify the setup of the plot grid.

Several grid types are available, with “Rectangle (polygon)” being best suited for many plant science applications. The user can then specify the total grid extent by drawing on the map canvas, and specify the horizontal and vertical spacing of the cells in the grid. Positive or negative overlays can be made so that polygon boundaries overlap or have buffered space between them, respectively. Negative (buffered) overlays, which only draw data from the central portions of each plot rather than the periphery, can be particularly helpful for certain data types. This is particularly true for thermal and vegetation index imagery, if elimination of edge effects is desired. An output destination should be selected to permanently save the grid as a geopackage.

Once the grid has been generated, it can be manipulated to ensure that it corresponds ideally with the actual field. If data is being captured on growing canopies throughout a time series, it is often best to start with a later timepoint, when plants have grown together or nearly so, and then use this grid as the basis for earlier timepoints, before canopy closure, when separation is simpler.

By default, the cells of the grid layer will be opaque, making it difficult to fine-tune the grid position. Transparency can be adjusted by right-clicking the layer in the Layers Panel to access the “Properties” settings, where in the “Symbology” tab there is an opacity adjuster. Similarly, to adjust the cells, they will need to be selected, and selected features are opaque yellow by default. This opacity setting for the project’s selected features can be changed in the Project Properties window (Project → Properties) under the “General” tab at left.

Many editing features are only available when the Advanced Digitizing Toolbar is activated (View → Toolbars → Advanced Digitizing Toolbar). The grid can be edited by then toggling editing using the pencil icon, while the grid layer is selected in the Layers Panel. Cells of interest can then be selected using the “Select features” tool in the toolbar. The cursor with the shift key held can be used to select groups of cells, which then can be edited. This can include deleting extra rows or columns, if present. The “Move” and “Rotate” functions may be particularly useful for adjusting the locations of polygons cells in the grid. Adjustments can be made by alternating between the select tools and move or rotate tools until the grid aligns well with the layers with data of interest.

In some cases, it may be valuable to extract data from the central part of each plot without edge effects. This can be done in the initial grid set up in the overlay sections (described previously), or a second grid can be developed which focuses specifically while maintaining extract data from the central part of each cell, using the Vector → Geoprocessing Tools → Buffer, then using negative values for the buffering distance.

When different sections of a field have different plot sizing or spacing, it can be helpful to set up multiple grids, and then combine them into a single merged file to simplify data extraction. This can be accomplished by Vector View → Toolbars → Advanced Digitizing Toolbar. The layers can then be selected and merged. The original vector file that each cell in the grid came from can be found in the attribute table of the new combined grid.

It is worth noting that after establishing a plot grid, the following steps including the Raster Calculator, Zonal Statistics, and data download, can be automated and run as a script in the Python Console. This is described in detail in section 2.11. However, the following sections are important for setting up the process, such as optimizing the values used for thresholds, and can serve as the basis of later automation. They also offer an intuitive graphical user interface, which may be preferred by many users, and may be easier to use for isolated flights. These methods will lay the groundwork for the automation methods of section 2.11.

### 2.5. Vegetation index calculation

Full protocol: https://plantsciencedronemethods.github.io/pages/6.VegetationIndices/Methods.html The Raster Calculator is an important function that will be the basis for the next several steps, including calculating vegetation indices. Vegetation indices are based on simple calculations of reflectances at different wavelengths. If photogrammetry was conducted in Pix4D, it is best to use index rasters from the 4_index folder, rather orthomosaics from the 3_dsm_mosaic folder. Once these are imported into QGIS, the Raster Calculator can be opened (Raster → Raster Calculator) and equations can be calculated by clicking on the layers of interest and using the mathematical operators or typing them with the keyboard in the Raster Calculator Expression box. Numerous vegetation indices are available and have been reviewed elsewhere (e.g., Xue and Su, 2017; Giovos, 2021). Among these, among the earliest and the most widely used (Huang et al., 2021) is the normalized difference vegetation index (NDVI), whose equation is the following:

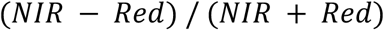

A permanent file path should be specified before running. After running, the data of individual points can be determined using the “Identify Features” tool in the toolbar, as long as the layer of interest is selected in the Layers Panel.

The appearance of vegetation index maps can be optimized by adjusting right-clicking on the layer of interest in the Layers Panel, then clicking Properties. In the Symbology tab, the Render Type can be changed to “Singleband pseudocolor”. Numerous color ramps are available, and scaling options are then available for editing.

### 2.6. Classification layers and masking soil in vegetation index or thermal files

Full protocols: https://plantsciencedronemethods.github.io/pages/8.ClassificationLayers/Methods.html

https://plantsciencedronemethods.github.io/pages/10.maskedNDVI/Methods.html

The Raster Calculator can be used to quickly and easily classify and distinguish plants and soil in the files of interest. This can be done using a simple raster calculator expression, such as:

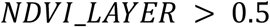

In this case, a value of 0.5 is used as the threshold for distinguishing plants from soil. This value may need to be adjusted for other vegetation indices, sensors, lighting conditions, and subjects.

The output of this raster calculation will be a binary classification layer. Pixels with NDVI values over 0.5 will be classified as plant, and will be stored as a “1” in the new layer. Pixels with NDVI values less than 0.5 will be considered non-plant, and will be saved as a “0” in the new classification layer.

The classification layer can be used directly to export canopy fraction and canopy cover, described later in the section on Zonal Statistics. The classification layer can also be used to quickly and easily remove soil pixels. For example, the following equation generates a masked vegetation index (NDVI) layer:

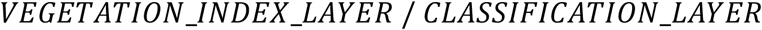

This will output a masked vegetation index (e.g., “mNDVI”) layer. In the classification layer, since soil pixels were stored as “0”s while canopy pixels were stored as “1”s, for canopy pixels this will return the same value as what was stored in the original NDVI file. In contrast, the division by 0 will result in “no data” for that pixel. This process eliminates the soil pixels. This is extremely important for the determination of canopy index layer information, without the confounding effects of soil when there is not full canopy closure. Similarly, soil pixels can be masked from thermal or other data, provided there is good alignment between files, using Raster Calculator expressions such as the following:

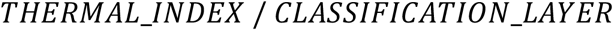

### 2.7. Height and volume calculations

Full protocol: https://plantsciencedronemethods.github.io/pages/11.HeightVolume/Methods.html

The Raster Calculator can also be applied to DSMs to extract data on the elevation of plant canopies and the surrounding soil. For example, to calculate the elevation of plant canopies, with soil masked, the following expression can be put into the Raster Calculator:

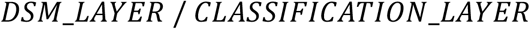

This will then output a canopy DSM (“cDSM”) file, that contains only elevation data from the plant canopies, with soil or non-plant features masked and stored as “no data” due to the division by 0. This is very helpful, as it allows for the examination of canopy height information, without the influence of soil elevation.

To accomplish a similar task, but to isolate soil heights while masking plant canopies, the equation can be updated and re-run:

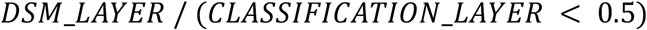

This will result in a soil DSM (“sDSM”). The expression “CLASSIFICATION_LAYER < 0.5” serves to functionally reverse the values of the classification layer: Non-soil pixels are stored as “0”s in the classification layer, which is less than the threshold of 0.5 in the expression, and will be saved as a “1” since the expression was true. For a canopy pixel, saved as a “1” in the classification layer, the expression is false, so that part becomes a “no data”. This eliminates canopy pixels without requiring additional files. These classification layers, mNDVI or other masked vegetation indices, cDSM, and sDSM files are then ready for data extraction using Zonal Statistics.

### 2.8. Data extraction: Canopy cover, vegetation indices, height, and volume using Zonal Statistics

Full protocol: https://plantsciencedronemethods.github.io/pages/9.CanopyArea/Methods.html

After generating masked rasters with data of interest, the data can conveniently be summarized and exported to attribute tables using the Zonal Statistics tool in QGIS. This interface allows for the selection of raster and vector input files, as well as the band in the raster image, and prefixes for the output columns, which may be the name of the raster file. The desired output statistics can be selected. Outputs are written to an attribute table, accessible by right-clicking the vector layer where data were output in the Layers Panel, then selecting “Open Attribute Table”. These data can be copied and pasted into other programs, such as Microsoft Excel, from which they can be saved and read into statistical software such as R.

The mean value of the classification layer for each plot gives the canopy fraction in that cell.

The total canopy area of the plots can be calculated by multiplying the mean value of the classification layer against the cell size, as specified in the “Create Grid” function earlier. Statistics such as masked vegetation indices and masked canopy temperature can be directly downloaded without further processing, although some thermal outputs may need to be converted to degrees Celsius based on a mathematical function (e.g., raw value × 0.04 - 273.15 = °C) provided by the camera manufacturer.

For determination of canopy height, the median value of the soil DSM will typically give the elevation of the base of the plants, while mean canopy DSM values represent the average height of plants. Values from Pix4D and Metashape will be given in meters. Mean and maximum plant canopy height can be calculated by subtracting median soil elevations (sDSM values) from each plot from the mean and maximum canopy height values, respectively. Volume can be calculated by multiplying the canopy area by the mean canopy height.

### 2.9. Splitting fields into plot-level imagery

Full protocol: https://plantsciencedronemethods.github.io/pages/11.5FieldSplit/Methods.html

In some cases, it may be useful to have images of individual plots, such as for running processes in ImageJ (e.g., stand counts, Sunoj et al., 2025), and for running computationally intensive image analysis processes. Once a plot grid has been generated (as described in section 2.4), the “Clip Raster by Mask Layer” function can be used to export images of the individual plots. In this function, the raster, vector grid layer, an output location, and base file name are required. It is critical to iterate over all plot cells, which is achieved by clicking the icon with green arrows in a circular format. The function will then output a series of images with the desired base name and location, which can be batch-processed in ImageJ or other software.

### 2.10. Raster Merging and Multispectral to True-Color Imagery

Full protocol: https://plantsciencedronemethods.github.io/pages/11.75MultiSpecToRGB/Methods.html

In some cases, it may be helpful to merge raster layers into single files. For example, the Red, Green, and Blue channels of multispectral cameras can be merged into a single file, which can then be used combined into a multiband raster to approximate a true-color RGB image. This can be done using the Raster → Miscellaneous → Merge function, and by selecting the multispectral red, green, and blue index rasters as input files. In the Properties → Symbology section of the new layer, the red and blue channels may need to be swapped, and the minimum and maximum values of each channel will need to be made consistent. The resulting output will be an RGB image based on multispectral index rasters.

Merging rasters can also save time in georeferencing. For example, multiple outputs from the same photogrammetry project can be merged into a single file, which then can be georeferenced a single time. DSMs can be particularly difficult to georeference on their own, due to the lack of 3-dimensionality of most GCPs, making the GCPs difficult to identify in the topographic models. The Merge function can be used to merge the layers of interest, such as these digital surface models, with other outputs from the same project, such as vegetation index maps. These can then be simultaneously georeferenced as one file, and data can be extracted directly from each individual band of the merged file.

### 2.11. Automating drone data extraction using the python console

Full protocol: https://plantsciencedronemethods.github.io/pages/12.AutomationScript/Methods.html

QGIS’s Python Console offers the opportunity to automate and simplify many workflows, particularly repetitive processing steps. This is extremely valuable when conducting repeated measurements of fields, or extracting similar data from different locations and dates. This automation can be applied for many parts of the drone data analysis pipeline, such as the calculation of vegetation indices, soil masking, and data extraction.

Python code can easily be downloaded for QGIS functions, when the functions are accessed from the Processing Toolbox. These are available under the “Advanced” option at the bottom of the function window. These can then be combined into longer workflows. By creating variables to store the names of files at the beginning of the script, and then referring to them in subsequent steps as the variable name, large data handling pipelines can be run by simply updating the names of input files at the beginning of the workflow.

### 2.12. Object detection, segmentation, and artificial intelligence in drone data processing

Artificial Intelligence (AI) based methods are a rapidly growing component of drone data processing pipelines. For example, the Orfeo Toolbox offers machine learning tools for image segmentation and a variety of classification features, which are particularly useful when working with multiple (3+) object classes in field imagery. These tools can be accessed by extracting the Orfeo Toolbox to the C:/ directory, and specifying the paths to the OTB application folder and OTB folder in Processing → Options → Providers → Orfeo Toolbox. To train a classification model, a vector layer can be created, and points or polygons can be specified and given specific numeric IDs corresponding to distinct classes for the model. This vector file can then be used as an input, along with its associated raster file, to train a classification model using the Orfeo Toolbox’s TrainImagesClassifier function. This model can in turn be used as an input for the ImageClassifier function of the Orfeo Toolbox, along with the raster file to generate a new classification layer based on the training data. The resulting output will include a new raster with each pixel classified into one of the category types, as well as a confidence map for each pixel.

Numerous other methods are available and rapidly expanding for the use of machine learning and computer vision in drone data processing. For example, feature annotation can be accomplished using Roboflow, which offers an intuitive user interface, but is proprietary software and processing large data sets or use of advanced tools requires a subscription. Alternatively, annotation can be accomplished in PyTorch or TensorFlow, which are free and open source and highly flexible, but may require greater user abilities in Python coding. Models can then be trained or selected from existing repositories. For example, the QGIS Deepness plugin (Aszkowski et al., 2023) includes a repository of existing models, as well as resources for custom model development using the Yolo model family (Redmon et al., 2016; Wang and Liao et al., 2024), although model training is done outside QGIS. The Deepness plugin allows for the implementation of deep neural networks to conduct segmentation, regression, and object detection on raster files, in a user-friendly interface from QGIS. Methods like these are an area of rapid growth and will continue to advance in the coming years.

## 3. Results and Discussion

### 3.1. Uses and validation

The methods described herein are applicable to the study of a wide variety of different traits across the plant kingdom (Figure 1). Generation and extraction of data using these methods is high-throughput, including the full automation of processing after photogrammetry and plot specification. Post-photogrammetric steps of this workflow can often be achieved in minutes. Critically, the calculated data are highly precise. For example, the correlation coefficient of vegetation indices can readily exceed R^2^ = 0.99, even when the vegetation indices were based on image sets from separate cameras mounted on separate aircraft and flying at different altitudes (Figure 1G). Similarly high correlations (R^2^ > 0.99) were achieved using methods described here when measuring canopy area from distinct cameras (Parker et al., 2020). These results highlight the high precision of the data derived from these methods. Strong correlations are also seen between numerous other related traits, such as canopy height, temperature, and a variety of other vegetation indices (Figure 1H-J).

**Figure 1.**
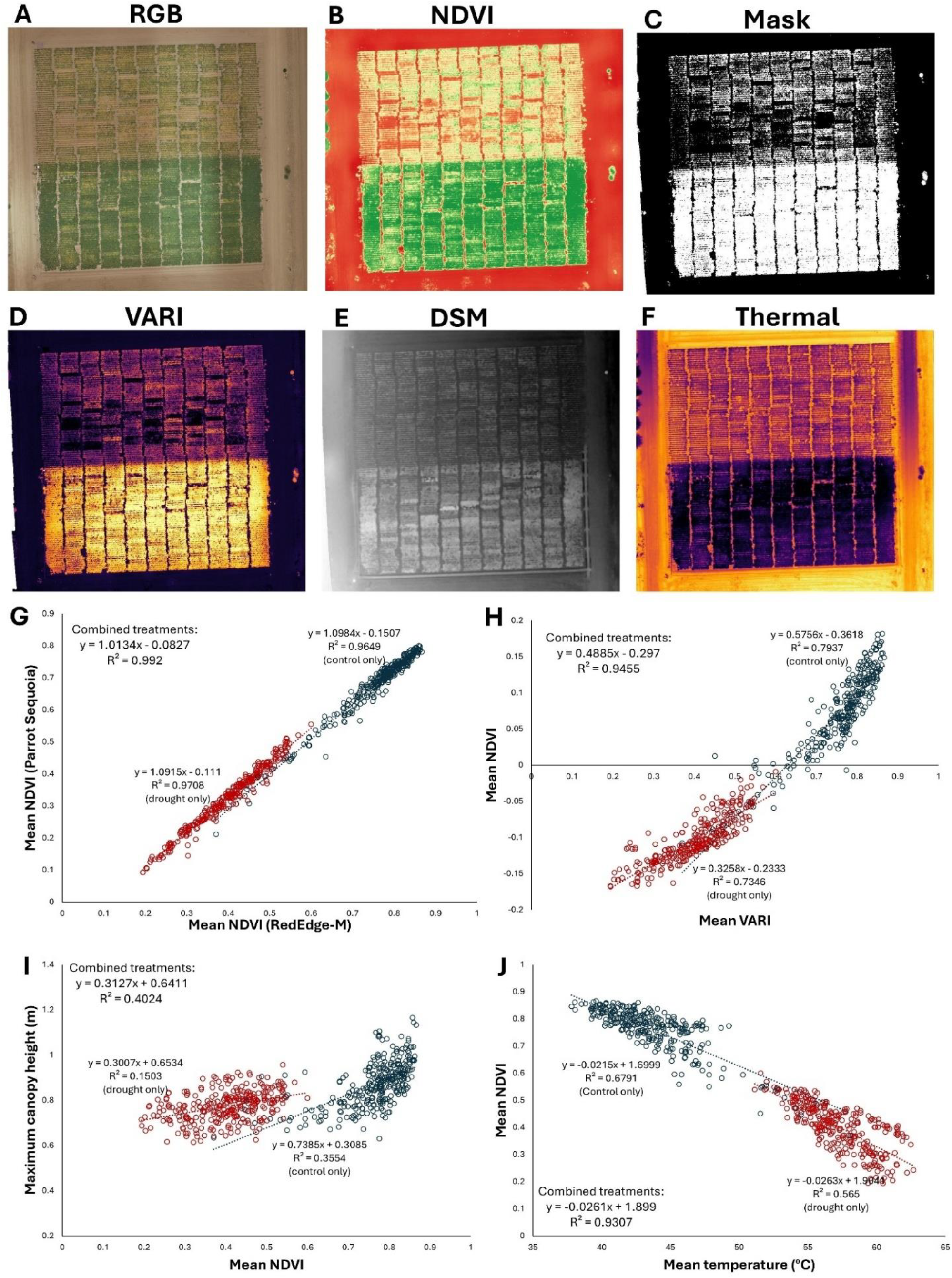
The methods described herein can be used to extract a variety of data types from many sensors, including multispectral, RGB, and thermal cameras. In this example, four cameras were flown over a terminal drought stress experiment in chickpea: (A) An RGB orthomosaic (drought stress at north, well watered control to the south) (B) NDVI, (C) NDVI-based classification layer delineating living plant material from non-living material, (D) visible atmospherically resistant index (VARI) (E) a digital surface model (DSM) topographic map, (F) thermal map of canopy temperature. (G) Comparison of NDVI values calculated over the field from distinct cameras, flown on separate aircraft at different altitudes, with data extracted from independently generated plot grids. The extremely strong correlation (R^2^ > 0.99) demonstrates the high precision and reliability of these data processing and extraction methods. (H) VARI from an inexpensive RGB stock camera is strongly correlated with NDVI from a multispectral Micasense RedEdge-M camera, (I) Correlation between canopy height and mean NDVI is comparatively low, indicating that the drought stress treatment strongly affects canopy NDVI but not canopy height, which was largely established before the onset of the terminal drought stress. (J) Canopy NDVI and temperature are strongly negatively correlated. All data processing conducted through methods described here and are previously unpublished. Field management by Kay Watt and Antonia Palkovic.

Subsets of these methods have been applied across numerous plant species. This includes the study of growth rate and canopy architecture genetics in beans and cover crops (Parker et al., 2020; Garcia Patlan, 2024), stem water potential in almond, pecan, and orchard trees (Savchik et al., 2025; Taylor et al., 2024), drought stress tolerance and response in beans (Wong et al., 2023; Buckley et al., 2025), disease detection in maize and wheat (Bevers et al., 2024; Buster et al., 2023), and soil analysis (Mei et al., 2023). Numerous other applications of these methods have also been conducted (Figure 2).

**Figure 2.**
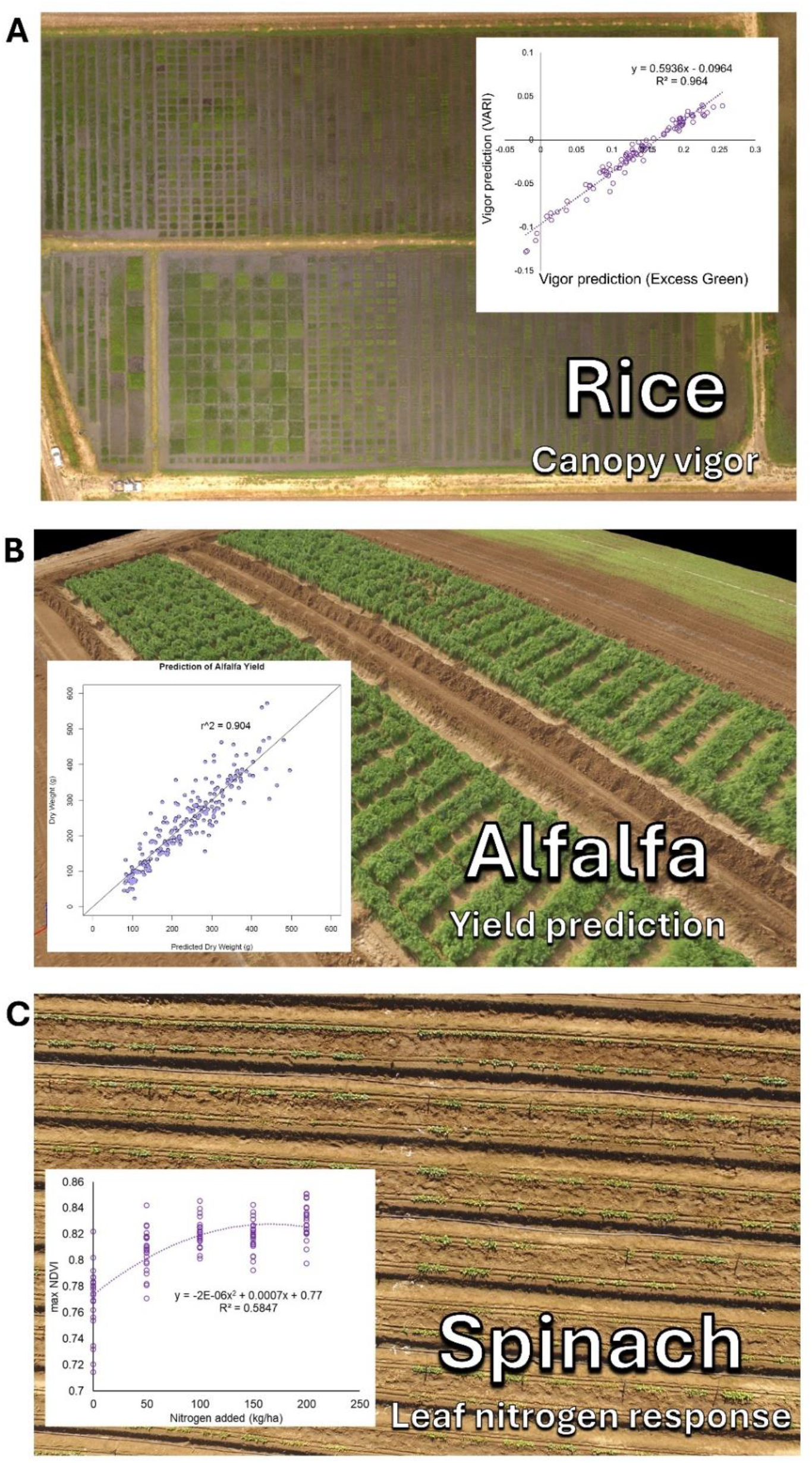
Examples of these methods applied to diverse plant science applications. (A) Canopy growth vigor characterization in rice breeding, (B) Yield prediction in alfalfa, and (C) Leaf nitrogen response in spinach.

### 3.2. Training resources

The methods described herein have been deposited in two main educational repositories, 1) a Plant Science Drone Methods website and 2) an educational YouTube channel (Figure 3). The web page, available at PlantScienceDroneMethods.github.io, features a series of step-by-step protocols for each part of the drone data analysis pipeline. The site also includes an example script for automated data processing and extraction, example data for users to test for their own analyses, and other resources such as relevant publications. The development of the Plant Science Drone Methods web page facilitates keeping methods protocols up to date with the latest technological advances and software updates.

**Figure 3.**
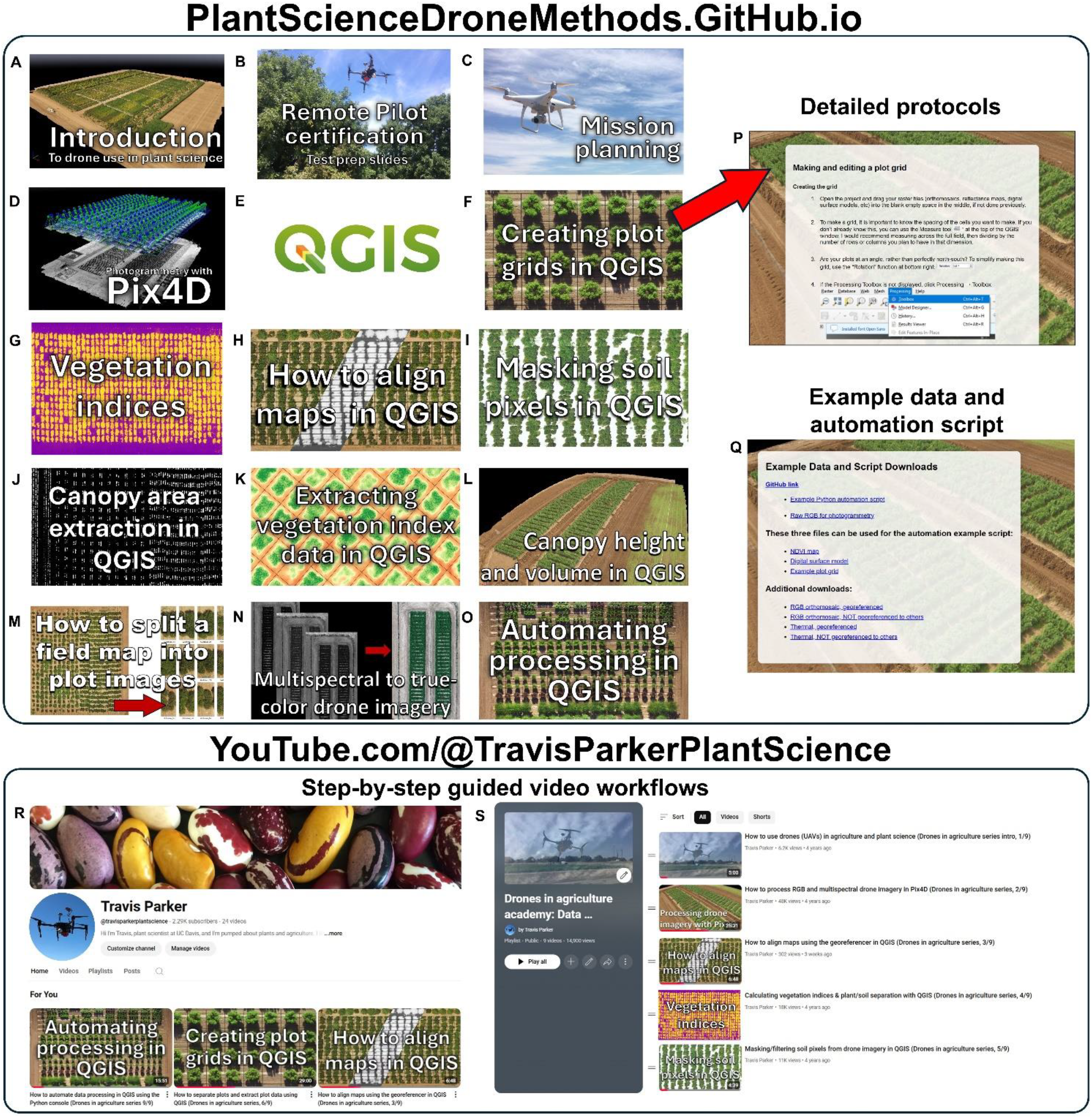
Educational and training resources for drone use in plant science. (A-O) Example protocol links on the Plant Science Drone Methods web page (PlantScienceDroneMethods.GitHub.io). Each links to (P) a detailed procedure. (Q) The web page also includes example data and an example automation script for users to try out the methods. (R) Additionally, an educational YouTube channel has been developed and hosts (S) a playlist covering the drone data generation and extraction pipeline. These serve as training resources in a particularly clear and thorough format. Both the Plant Science Drone Methods page and YouTube channel are freely available and focus primarily on free- and open-source software, promoting global accessibility.

Similarly, step-by-step instructional videos have been published on an educational YouTube channel and playlist (https://www.youtube.com/watch?v=obRng3LLkRs&list=PLifbxiLZAb-QHX2QjBqCNKysSHM3gPuTX). These methods help demonstrate complex software steps in a clear and transparent way, breaking down both the theory and practice of each step of the data extraction pipeline. They also serve as a discussion forum for asking questions and giving comments on uses.

These free online resources focus primarily on free and open-source options. This promotes their accessibility to communities that have traditionally lacked access to expensive proprietary software or centers of higher learning. The development of these freely available resources has and will continue to unlock the potential of drones to serve as platforms for the collection of high-precision, high-throughput data in plant science.

## 4. Conclusion

Drones have enabled the capture of high-resolution data for a variety of plant traits. Here, we describe methods and educational resources required to extract meaningful plant canopy data from these emerging technologies. These steps include the processing of raw images into photogrammetric 2D and 3D outputs, georeferencing and aligning them between flight dates, generating plant vs. soil classification layers, calculating vegetation indices based on reflectance data, specifying plot boundaries by generating plot grid geopackage layers, extracting the data into tabular format, and using machine-learning approaches to annotate and classify drone-derived images. These tools are flexible and can be combined and interwoven to study a diverse range of plant traits. These methods and resources will facilitate the high-throughput phenomic characterization of large plant populations, ultimately improving the productivity, sustainability, and ecological health of plant populations globally.

## 6. Author Contributions

Conceptualization: TAP, PG; Data curation: TAP; Funding acquisition: TAP, PG; Methodology: TAP, BC, MW; Software: TAP, BC, MW; Writing – original draft: TAP; Writing – review & editing: TAP, PG, BC, MW.

## 8. Acknowledgments

Taylor Nelsen and Alex Mandel provided extremely valuable help with QGIS. Antonia Palkovic, Kay Watt, Charlie Brummer, Matt Francis, Scott Newell, Oon Ha Shin, and Allen Van Deynze contributed to field preparation and management. Brandon Stark provided help with hardware and regulatory questions.

## 1 Data Availability Statement

Data and code can be found at PlantScienceDroneMethods.GitHub.io

